# The origin of mouse extraembryonic endoderm stem cell lines

**DOI:** 10.1101/2021.12.27.474300

**Authors:** Jiangwei Lin

**Author notes:** J. Lin: Kunming Institute of Zoology, Kunming, Yunnan 650223, China. Correspondence: Jiangwei Lin.

## Abstract

Mouse extraembryonic endoderm stem (XEN) cell lines can be derived from preimplantation embryos (pre-XEN) and postimplantation embryos (post-XEN). XEN cells share a gene expression profile and cell lineage potential with primitive endoderm (PrE) blastocysts. However, the cellular origin of XEN cells in embryos remains unclear. Here, we report that post-XEN cell lines are derived both from the extraembryonic endoderm and epiblasts of postimplantation embryos and that pre-XEN cell lines are derived both from PrE and epiblasts of blastocysts. Our strategy consisted of deriving post-XEN cells from clumps of epiblasts, parietal endoderm (PE) and visceral endoderm (VE) and deriving pre-XEN cell lines from single PrE and single epiblasts of blastocysts. Thus, XEN cell lines in the mouse embryo originate not only from PrE and PrE-derived lineages but also from epiblast and epiblast-derived lineages of blastocysts and postimplantation embryos.

## INTRODUCTION

After the 8- to 16-cell stage, mouse embryos start to compact and initiate the first cell fate decision to form two distinct cell types: the outer and inner cells. The outer cells will develop mostly into trophectoderm, which ultimately give rise to the fetal portion of the placenta. The inner cells will develop mostly into the inner cell mass (ICM) ^1^. After the 32- to 64-cell stage and the formation of the early blastocyst, embryos start the second cell fate decision: ICM develops into epiblast and PrE. The epiblast gives rise to most of the embryo proper, amnion, and extraembryonic mesoderm of the yolk sac. PrE forms the extraembryonic endoderm (ExEn) lineage: visceral endoderm (VE) and parietal endoderm (PE) of the yolk sac^2-4^.

Various types of stem cell lines can be derived from preimplantation and postimplantation embryos. Embryonic stem (ES) cell lines are derived from blastocysts ^5,6^ and can be converted from postimplantation embryos ^7^. Trophoblast stem (TS) cell lines are derived from blastocysts and extraembryonic ectoderm (EXE) of postimplantation embryos ^8-10^. Extraembryonic endoderm stem (XEN) cell lines were first derived from blastocysts.^11^ XEN cell lines can also be converted from ES cell lines by ectopic gene expression or by adding chemical factors to the culture medium^12-17^ or be induced from fibroblasts ^18^. We have previously reported that XEN cell lines can be derived efficiently from postimplantation embryos ^19^ and that the derivation and maintenance of XEN cell lines does not require PDGRFA^20^. We termed XEN cell lines derived from postimplantation embryos “post-XEN cell lines” to distinguish them operationally from XEN cell lines derived from blastocysts, which we termed “pre-XEN cell lines”. The embryonic origin of pre-XEN and post-XEN cell lines remains unclear. Mouse fibroblasts pass through an XEN-like state before forming induced pluripotent stem cells by chemical reprogramming^21^.

Here, we derived post-XEN cell lines from clumps of epiblasts, PE and VE and derived pre-XEN cell lines from single PrE and single epiblasts of blastocysts.

## RESULTS

### Derivation of post-XEN cell lines from E6.5 (R26-tauGFP41 x Sox17-Cre) F1 embryos

We set up a natural mating between a homozygous female of the gene-targeted Cre reporter strain R26-tauGFP41^22^ and a heterozygous male of the gene-targeted driver strain Sox17-Cre^23^. In (R26-tauGFP41 x Sox17-Cre) F1 embryos, cells become permanently labeled with GFP upon expression of Sox17 (which occurs in PrE but not in epiblasts), and the descendants of these cells, including XEN cells, are also labeled permanently.

We have reported the derivation of post-XEN cell lines from postimplantation embryos at E6.5 ^19,20^. We sectioned 14 (R26-tauGFP41 x Sox17-Cre) F1 decidua at the E6.5 stage and found that the overwhelming majority of GFP+ cells resided within the ExEn (Fig. 1A). Then, we collected 5 GFP+ and 4 GFP-E6.5 (R26-tauGFP41 x Sox17-Cre) F1 embryos from a natural mating between a homozygous R26-tauGFP41 female and a heterozygous Sox17-Cre male. Previously, we derived post-XEN cell lines from postimplantation embryos at E6.5 with whole embryos (termed the whole embryo method) ^19,20^. Here, we incubated the embryos in 2 mg/ml collagenase in Ca^2+^/Mg^2+^-free PBS for 20 min at room temperature and disaggregated them into small pieces and single cells using a 120-200 *µ*m glass pipette (termed the disaggregated embryo method) (Fig. 1B). We placed each disaggregated embryo into a well of a 4-well dish in TS cell medium with F4H. The cultures of the GFP+ embryos were intended to establish post-XEN cell lines, and the cultures of the GFP-embryos were used for immunofluorescence analysis. On day 4, we observed two types of colonies: GFP+ XEN-like colonies and GFP-flat colonies (Fig. 1C). We picked colonies of the two types using two hypodermic 30G needles and a 20-*µ*l plastic pipette tip and pooled each colony type. We disaggregated each pool into a suspension of small pieces and single cells and transferred the suspension into a well of a 4-well dish in TS cell medium with F4H. Cultures from pooled GFP+ XEN-like colonies grew quickly, and we established 5 post-XEN cell lines from 5 pools (Fig. 1D). Cultures from pooled GFP-flat colonies initially maintained a flat morphology, with GFP+ cells surrounding on day 10; then, on day 35, only GFP+ XEN-like cells remained (Fig. 1E). It appears that cells of GFP-flat colonies convert into GFP+ XEN-like cells in culture. We established 3 post-XEN cell lines from 5 pools of GFP-flat colonies.

**Figure 1.**
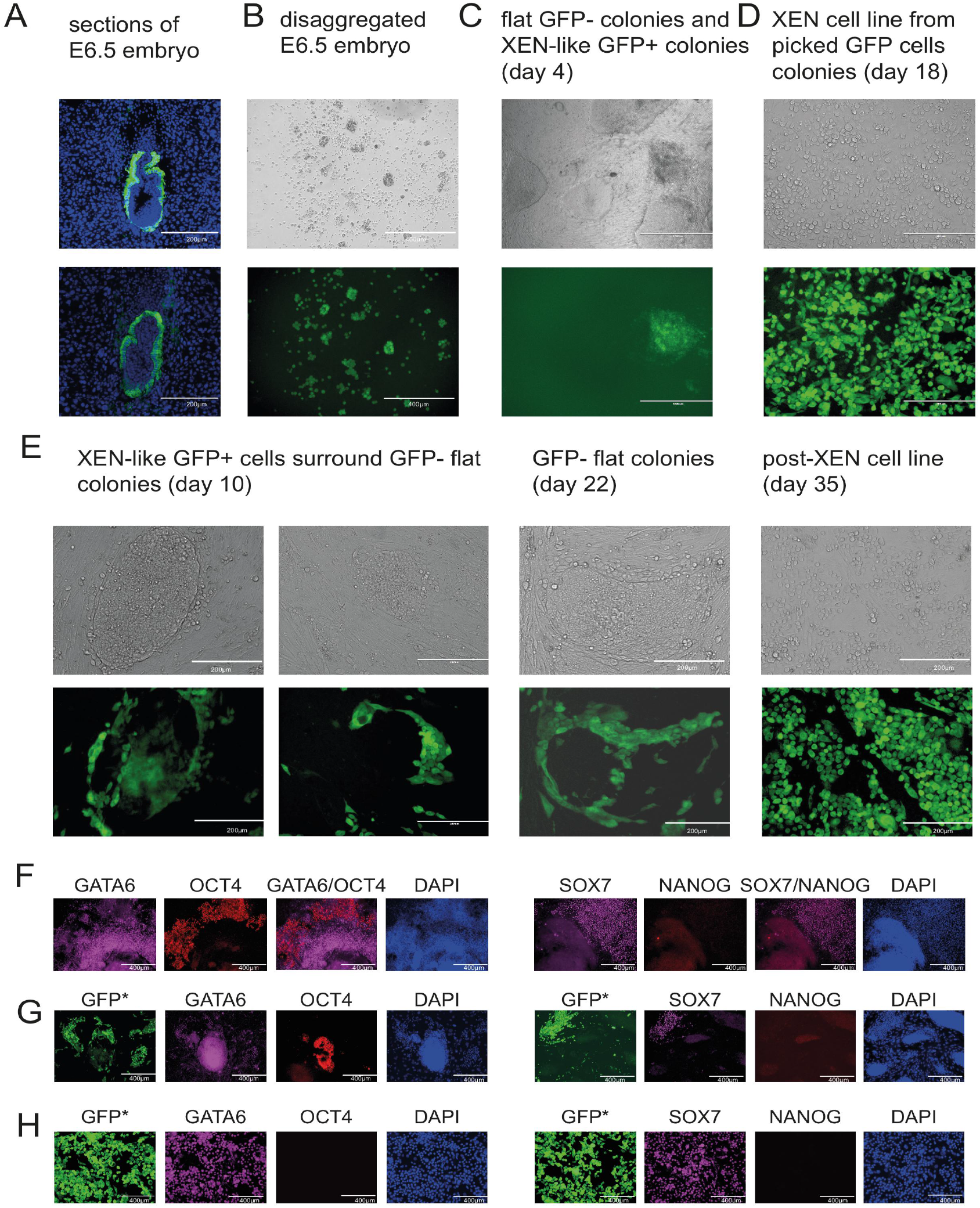
Derivation of post-XEN cell lines in E6.5 (R26-tauGFP41 x Sox17-Cre) F1 embryos. (**A**) Cryosections of two E6.5 (R26-tauGFP41 x Sox17-Cre) F1 deciduae. Thickness is 12 *µ*m. Intrinsic green fluorescence of GFP, counterstaining with DAPI (blue). (**B**) E6.5 embryo without ectoplacental cone and same embryo after disaggregation. Top: bright-field image; bottom, intrinsic green fluorescence of GFP. (**C**) There were two types of colonies on day 4: flat and GFP- and XEN-like and GFP+. (**D**) Post-XEN cell line X-E6.5-AC1456-4 on day 18, derived from a pool of XEN-like colonies picked on day 5. Most cells in the culture are GFP+. (**E**) GFP+ XEN-like cells (X-E6.5-AC1456-4) surrounding GFP-flat colonies on days 10 and 22. On day 35 there were no more GFP-flat colonies. (**F**) Immunofluorescence was performed on outgrowths from GFP-E6.5 embryos on day 7. (**G**) Flat colonies displayed GATA6 and SOX7 expression, strong OCT4 expression, and weak or no NANOG expression on day 18. (**H**) Most cells from flat colonies had the typical appearance of XEN-like cells, with strong expression of GATA6 and SOX7 and no OCT4 or NANOG expression on day 36.

We performed immunofluorescence on outgrowths from disaggregated GFP-E6.5 embryos on day 7 and on cultures on days 18 and 36. The outgrowths expressed GATA6, SOX7 and OCT4 and weakly or no NANOG on day 7 (Fig. 1F). On day 18, flat colonies displayed GATA6 and SOX7 expression, strong OCT4 expression, and weak or no NANOG expression (Fig. 1G). Marker expression in flat colonies distinguishes them from epiblast stem cells (which express NANOG and GATA6) and ES cells (which express NANOG but not GATA6)^24-26^. On day 36, most cells from flat colonies had the typical appearance of XEN-like cells, with strong expression of GATA6 and SOX7 and no OCT4 or NANOG expression (Fig. 1H).

### Tracing the origin of post-XEN cell lines in E6.5 (R26-tauGFP41 x Sox17-Cre) F1 embryos

Next, to trace the origin of XEN cells from postimplantation embryos, it is better to isolate single cells from ExEn and epiblasts of postimplantation embryos to derive XEN cell lines. Unfortunately, we never obtained a post-XEN cell line from a single cell cultured in a 4-well dish and 96 plate in TS medium with F4H and never obtained a post-XEN cell line from all single cell suspensions of an embryo. It could be that making ExEn and epiblast cells to single cells would damage the cells. Another reason could be that single ExEn and single epiblast cells could not grow well in the single-cell stage. A previous study reported that epiblast stem cells would be induced widespread cell death after being disaggregated to single cells by trypsin or other single-cell dissociation methods^24^. However, clumps from disaggregated methods could survive and establish the post-XEN cell line ^19^. We traced the origin of XEN cells in E6.5 (R26-tauGFP41 x Sox17-Cre) F1 embryos in clumps from ExEn, epiblasts and EXE. In (R26-tauGFP41 x Sox17-Cre) F1 embryos, cells become permanently labeled with GFP upon expression of Sox17 (which occurs in PrE but not in epiblasts), and the descendants of these cells, including XEN cells, are also labeled permanently ^23^. GFP+ cells were located in ExEn, and epiblasts were located in the middle of the embryo piece and consisted of GFP-cells. We produced 4 GFP+ E6.5 (R26-tauGFP41 x Sox17-Cre) F1 embryos by natural mating between a homozygous R26-tauGFP41 female and a heterozygous Sox17-Cre male (Fig. 2A). We used two hypodermic 30G needles to cut each embryo into three pieces: an extraembryonic ectoderm piece (EXE piece) consisting of EXE and ExEn surrounding EXE; an epiblast piece (EPI piece) consisting of epiblast and ExEn surrounding epiblast; and a middle piece consisting of the transition between EXE and epiblast (Fig. 2B). We discarded the middle piece. We incubated the EXE and EPI pieces separately in 2 mg/ml collagenase in Ca^2+^/Mg^2+^-free PBS for 20 min at room temperature. First, we used a glass needle of 200 *µ*m inner diameter to separate EXE, EPI, and ExEn pieces. Using a glass needle of 60-80 *µ*m inner diameter, we disaggregated the EXE, EPI and ExEn pieces into small clumps. We then picked several clumps of ExEn (combining ExEn from EXE and EPI pieces), EPI, and EXE. The ExEn clumps expressed GFP and had a dark background; in contrast, the EPI and EXE clumps did not express GFP and had a white background (Fig. 2C) because any GFP+ cells in the EPI and EXE clumps were detected by fluorescence on day 0. We pooled and transferred each of the three types of clumps into a well of a 4-well dish in TS cell medium with F4H. On day 2, the ExEn clumps showed strong GFP fluorescence; the EPI clumps showed weak GFP fluorescence (on day 0, no GFP fluorescence); and the EXE clumps had no GFP fluorescence. On days 4 and 6, the three types of clumps formed outgrowths. On day 16, the ExEn outgrowths abounded with GFP+ XEN-like cells; the EPI outgrowths were a mixture of GFP+ XEN-like cells and GFP-cells; and the EXE outgrowths were devoid of GFP+ cells. Thus, GFP+ cells from E6.5 (R26-tauGFP41 x Sox17-Cre) F1 embryos could originate from ExEn and can also be converted in culture from epiblasts but not from EXE.

**Figure 2.**
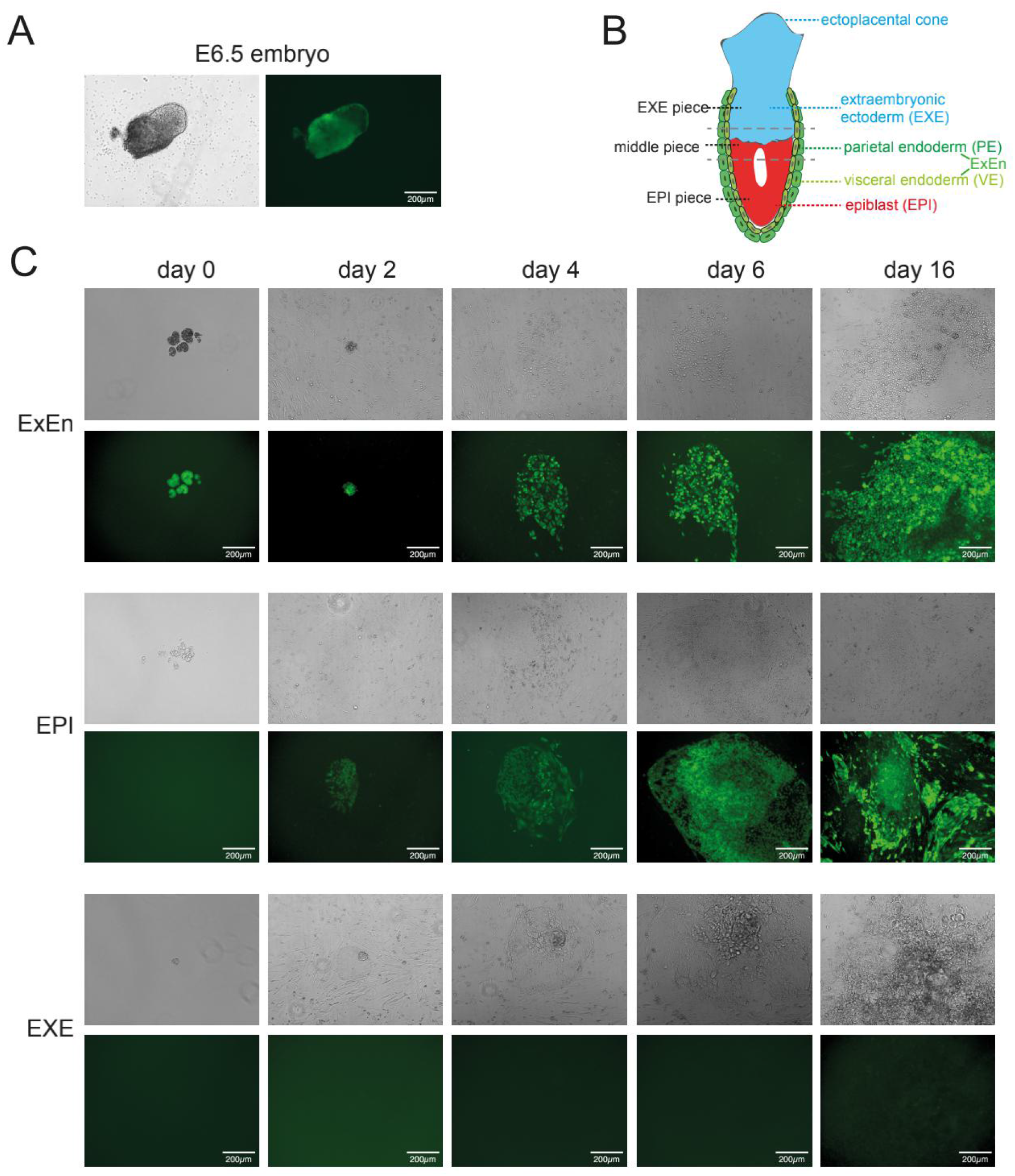
Tracing the origin of post-XEN cells in E6.5 (R26-tauGFP41 x Sox17-Cre) F1 embryos. (**A**) E6.5 embryo. Left, bright-field image; right, intrinsic green fluorescence of GFP. (**B**) Schematic representation of an E6.5 mouse embryo. The EXE piece contains EXE and ExEn; the middle piece, which is discarded, contains some EXE and some EPI; and the EPI piece contains EPI and ExEn. (**C**) Culture of clumps of ExEn, EPI, and EXE on various days.

### Tracing the origin of post-XEN cell lines in E7.5 (R26-tauGFP41 x Sox2-Cre) F1 embryos

We have already derived post-XEN cell lines from E5.5 and E6.5 embryos ^19^. Here, we derived post-XEN cell lines from E7.5 embryos. We produced 4 GFP+ E7.5 (R26-tauGFP41 x Sox17-Cre) F1 embryos by natural mating between a homozygous R26-tauGFP41 female and a heterozygous Sox17-Cre male. By the disaggregation method, we picked several clumps of ExEn and as few epiblast cells as possible from each embryo into a well of 4-well dishes coated with gelatin, covered them with MEF, and cultured them in TS medium with F4H. On day 7, we picked XEN-like colonies, and on day 30, we established 4 post-XEN cell lines from 4 E7.5 embryos (Fig. 3A). To confirm that post-XEN cells from E7.5 can contribute to ExEn, we injected two post-XEN cell lines (X-E7.5-AC1563-1 and X-E7.5-AC1563-2) into 17 C57BL/6J blastocysts and transferred 17 blastocysts into two foster mothers. We obtained 13 deciduae of E7.5 stage, sectioned the deciduae, and then stained the sections with an antibody against PDGFRA and E-cadherin. We found that 6 chimeras from 13 deciduae with GFP cells contributed to ExEn (Fig. 3B).

**Figure 3.**
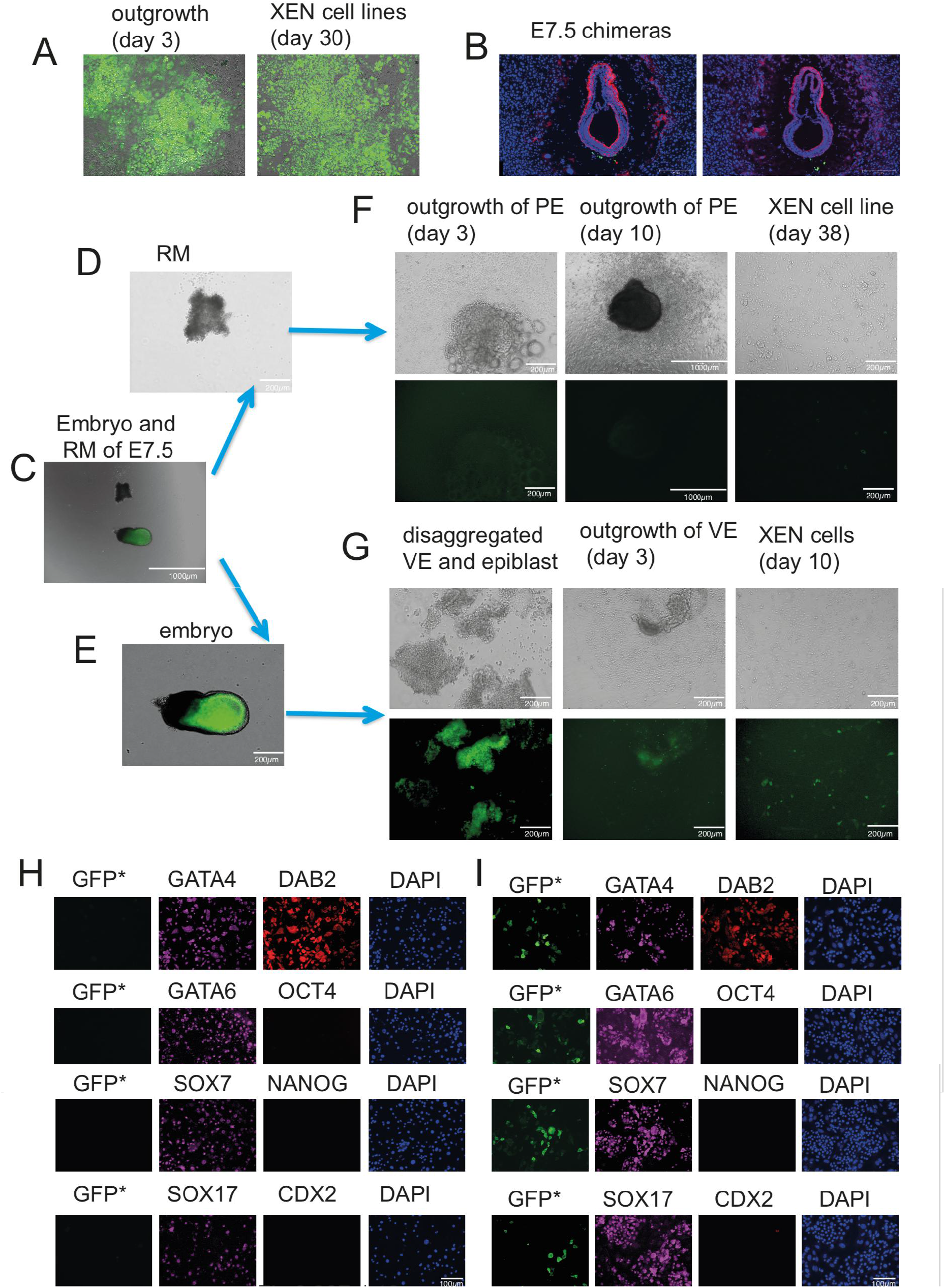
Tracing the origin of post-XEN cell lines in E7.5 (R26-tauGFP41 x Sox2-Cre) F1 embryos. (**A**) outgrowths from disaggregated E7.5 embryo. (**B**) Post-XEN cells contributed to ExEn in E7.5 stage embryos. (**C**) E7.5 embryo (bottom) and Reichert’s membrane (RM) (top). The bright-field image was merged with the fluorescence image. (**D**) RM. (**E**) Embryo without RM. (**F**) Outgrowth of RM (PE) on days 3 and 10 and post-XEN cell line X-E7.5-PE-Z1544-1 on day 38. All cells were GFP-. Top: bright-field image; bottom, intrinsic green fluorescence of GFP. (**G**) VE (GFP-) and epiblast (GFP+); outgrowth of VE on day 3; a small number of XEN cells on day 10 were GFP+. Top: bright-field image; bottom, intrinsic green fluorescence of GFP. (**H, I**) Post-XEN cell lines X-E7.5-PE-Z1544-5 (**H**) and X-E7.5-VE-Z01966-1 (**I**). Intrinsic green fluorescence of GFP and immunofluorescence for GATA4, GATA6, SOX7, SOX17, DAB2, OCT4, NANOG, and CDX2. DAPI was used as nuclear strain.

The transgenic driver strain Sox2-Cre is epiblast-specific^27^ and has been widely used to mark epiblast-derived cell lineages. Sox2-Cre strain is heterozygous^27^. Finally, to determine whether post-XEN cell lines are derived from PE and/or VE, we separated 11 GFP+ E7.5 (R26-tauGFP41 x Sox2-Cre) F1 embryos (Fig. 3C) into two pieces: Reichert’s membrane (RM) (Fig. 3D) and an embryo piece (epiblast with VE and EXE) (Fig. 3E). RM is formed by PE cells together with trophoblast giant cells^28^.

GFP+ cells were located in the epiblast region, which is in the middle of the embryo piece. VE surrounds epiblasts and EXEs and consists of GFP-cells. We cut the embryo piece into two pieces (EXE piece and EPI piece) with two hypodermic 30G needles and discarded the EXE piece, which was GFP-. We transferred a piece of each RM and each EPI piece (epiblast with VE) separately into a well of a 4-well dish in TS cell medium with F4H. On days 3 and 10, an outgrowth of XEN-like cells surrounded the RM, and 7 of 11 did not contain GFP+ cells, but 4 of 11 contained a few GFP+ cells (Fig. 3F). On day 12, we disaggregated the outgrowths and passaged the cells. On day 38, we established 9 post-XEN cell lines from 11 RM pieces (called PE-XEN). We treated 6 of the 11 EPI pieces with 2 mg/ml collagenase in Ca^2+^/Mg^2+^-free PBS for 10-20 min at room temperature and disaggregated them into small clumps using a 200-300 *µ*m glass pipette (Fig. 3G). We picked the VE clumps and cultured them in a well of a 4-well dish in TS cell medium with F4H. The VE clumps formed large outgrowths on day 3. Between days 3 and 7, we disaggregated the outgrowths into single cells and smaller clumps and passaged cells into a well of a 4-well dish in TS cell medium with F4H. On day 10, most cells in cultures from pooled VE clumps were GFP-. We established 6 post-XEN cell lines from the 6 disaggregated VE outgrowths (called VE-XEN). The other five EPI pieces were not treated with collagenase and were not disaggregated and did not give rise to post-XEN cell lines. Figure 3H shows immunofluorescence staining of the PE-XEN cell line X-E7.5-PE-Z1544-5 and Figure 3I of the VE-XEN cell line X-E7.5-VE-Z01966-1. These cell lines are immunoreactive for GATA4, GATA6, SOX7, SOX17, and DAB2 but negative for OCT4, NANOG, and CDX2. Thus, post-XEN cell lines of E7.5 embryos can be derived from PE and VE.

### Pre-XEN cell lines are both from single PrE and single epiblasts of blastocysts

To investigate whether pre-XEN cell lines originate both from PrE and epiblasts, we derived pre-XEN cell lines from single PrE cells and single epiblast cells. We set up natural matings between two homozygous R26-tauGFP41 females and two hemizygous Sox2-Cre males. We chose Sox2-cre instead of Sox17-cre because if epiblasts convert to XEN cells that would express Sox17 and then all cells would be GFP, at least for Sox2-Cre as a control, it is easier to judge epiblast-derived outgrowth in addition to the phenotype. In (R26-tauGFP41 x Sox2-Cre) F1 embryos, because GFP has not been expressed in the E4.5 blastocyst stage, single PrE and epiblasts can be distinguished until clones with a typical epiblast or PrE appearance have been formed: epiblast clones strongly express GFP and have an ES-like clone phenotype, and PrE clones do not express GFP and have an XEN-like clone phenotype.

We obtained 18 ICM immunosurgically^29^ from 18 (R26-tauGFP41 x Sox2-Cre) F1 E4.5 blastocysts from two females. We treated 15 of the 18 ICMs with TrypLE Express for 15 min in a CO_2_ incubator and then isolated single cells from every ICM with a 30 *µ*m glass pipette. ICMs could have approximately 20 single ICM cells^30^, and all single cells from ICMs were used as a group (Fig. 4A). If the group has any outgrowth cells that express GFP+, we term the ICMs R26-tauGFP41 x Sox2-Cre. If the group had no outgrowth cells that expressed GFP+, the ICMs were considered wild-type sox2. We placed 170 single cells from the 15 ICMs each into a well of a 4-well dish (all single cells did not express GFP) and cultured them in ES cell medium with LIF.

**Figure 4.**
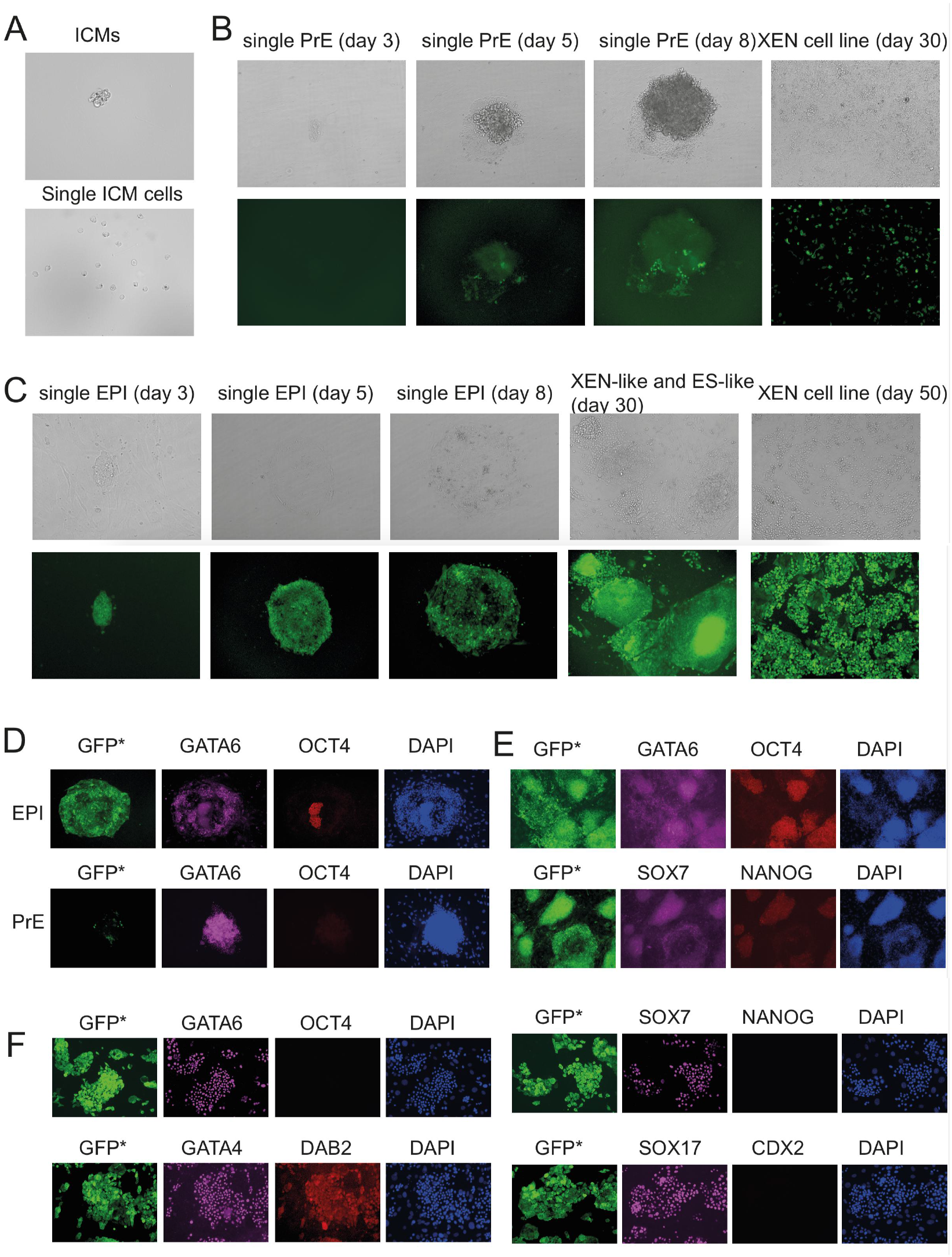
Pre-XEN cell lines are both from single PrE and single epiblasts of blastocysts. (A) Isolated single cells from an ICMs. (B) A single PrE formed outgrowth that was GFP- and with the phenotype of XEN-like on day 3, with a few GFP+ cells on day 5, day 8 and day 30. (C) A single epiblast formed outgrowth that was strongly GFP+ expression with an ES-like phenotype on day 3, day 5, day 8, and day 30 and converted to the XEN cell line on day 50. (D) Performed outgrowths could be derived from single epiblast cells from GFP+ ICMs for immunofluorescence on day 7. Performed outgrowths with few or no GFP cells and with an XEN-like phenotype could be derived from single PrE cells for immunofluorescence on day 7. (E) Immunofluorescence staining of ES-AC1558-2-8 from a single epiblast. (F) Immunofluorescence staining of cXEN-AC1558-2-8 converted from ES-AC1558-2-8 on day 61. DAB2, GATA4, GATA6, SOX17 and SOX7 were expressed in a single epiblast cell-derived XEN cell line. OCT4, NANGO and CDX2 were no expression.

We observed two different outgrowths from single ICM cells from days 3 to 8. The outgrowth with GFP-expression and a phenotype of XEN-like on day 3 was termed a single PrE, and the outgrowth with strong GFP+ expression and an ES-like phenotype on day 3 was termed a single epiblast (Fig. 4B-4C). However, unexpectedly, on days 5 and 8, we observed that a few GFP+ cells appeared in the outgrowths that were previously termed single PrE cells, and on day 30, a few GFP+ cells were still in the pre-XEN cell line, which was established from a single PrE (Fig. 4B). This result indicated that the Sox2-cre driver was activated in a few PrE cells during the XEN cell line derivation procedure. The outgrowths from single epiblast cells showed typical ES-like cells on day 3, day 5, and day 8. After disaggregating the outgrowth to single cells and clumps and passaging the cells, both ES-like and XEN-like colonies appeared in the plates (Fig. 4C). We got ES-like cells with some XEN-like cells. We cultured the cell lines either in ES medium with LIF or in TS medium with F4H on day 30. We changed the medium every other day and passaged the cells every 1-2 weeks. The cells were cultured with crowded, and some of XEN-like cells were in suspension medium, we collected the suspension medium and spun down, and passaged the cells into a new plate which coated with gelatin and covered with MEF, according to the protocol as we shown in previous report^20^, but we didn’t treat the cells with retinoic acid (RA) and activin A. We got XEN cell lines by collected the cells in suspension medium, in TS medium with F4H or in ES medium with LIF.

To confirm outgrowth with a typical XEN-like phenotype and that most cells were GFP-, which was termed from a single PrE cell, or outgrowth with a typical ES-like phenotype, and that most cells were GFP+, which was termed from single EPI cells, we performed immunofluorescence for the outgrowths that were from the ICMs of R26-tauGFP41 x Sox2-Cre on day 7. Four of 8 outgrowths had GFP+ expression and an ES-like phenotype. Some cells that were in the middle of the outgrowths expressed OCT4+, but GATA6+ expression was very weak. Therefore, these outgrowths could be derived from single epiblast cells. Another 4 of 8 outgrowths that had little or no GFP cells and with an XEN-like phenotype had strong GATA6+ expression and no OCT4+ expression; therefore, we concluded that these outgrowths could be derived from single PrE cells. Figure 4D shows immunofluorescence staining of such epiblast-derived outgrowth and PrE-derived outgrowth on day 7. In epiblast-derived outgrowth, cells were GFP+, some cells that were in the middle of the outgrowth still expressed OCT4+ and surrounded by GATA6+ cells, and OCT4 cells coexpressed GATA6, while the expression was very weak. PrE-derived outgrowths had GATA6 expression but no OCT4 expression, and there were a few GFP+ cells that arose from single PrE cells (Fig. 4D). Thus, by performing immunofluorescence, we confirmed that Pre-XEN cells could be derived both from a single PrE and from a single epiblast.

Figure 4E shows immunofluorescence staining of ES-AC1558-2-8, which was from a single epiblast, cultured crowded in ES medium with LIF on day 15. Which shows two distinct cell types: one is ES-like cells that is with OCT4 and NANOG expression, some of the ES-like cells are with weak GATA6 and SOX7 expression; another one is XEN-like cells that are with GATA6 and SOX7 expression. Figure 4F shows immunofluorescence staining of cXEN-AC1558-2-8 that converted from ES-AC1558-2-8 on day 61. DAB2, GATA4, GATA6, SOX17 and SOX7 are expressed in a single epiblast cell-derived XEN cell line, while OCT4, NANGO and CDX2 are not expressed.

Finally, we confirmed that out of the 15 ICMs, 6 ICMs were R26-tauGFP41 x Sox2-Cre. We isolated 73 single cells from the 6 ICMs. From the 73 single cells, we obtained 31 outgrowths, of which 18 outgrowths were from single PrE cells and 13 outgrowths were from single epiblast cells. Fifteen out of the 18 outgrowths had a few GFP+ cells, and 3 out of the 18 outgrowths had no GFP+ cells. In total, we obtained 6 pre-XEN cell lines from 14 single PrE cell-derived outgrowths (4 of the 18 outgrowths had already undergone immunofluorescence) and 4 pre-XEN cell lines from 9 single epiblast cell-derived outgrowths (4 of the 13 outgrowths had already undergone immunofluorescence). Thus, deriving pre-XEN cell lines from single ICM cells clearly shows that pre-XEN cells are derived not only from PrE but also from epiblasts.

## DISCUSSION

Typically, mouse XEN cell lines are derived from blastocysts^31^. We have recently reported that XEN cell lines can be derived efficiently from postimplantation embryos ^19,20^. The origin of XEN cells in blastocysts and in postimplantation embryos remains unclear. As XEN cells share a gene expression profile and cell lineage potential with PrE, it is traditionally assumed that XEN cells are derived from PrE^11^. XEN cells can be differentiated directly from mouse ES cells using growth factor^12^, and ES cells retain the ability to give rise to XEN-like cells spontaneously from the minor population expressing PDGFRA^32^. It would therefore be not surprising that conversion of epiblast-derived cells into XEN cells may occur during the derivation process from mouse preimplantation embryos. However, the establishment of XEN cells from the epiblasts of postimplantation embryos is a novel and interesting phenomenon. ES cell lines can be derived from earlier postimplantation embryos^7,^ and epiblast stem cells were reported to fail to obtain the XEN cell line^12^, suggesting that this conversion could occur via an ES-like intermediate state rather than an epiblast stem-like intermediate state. Anther thing is, thus far no paper has proved that pre-XEN cells are both from PrE and epiblast of blastocyst. Here, we have shown that post-XEN cell lines are derived from ExEn and epiblasts and that pre-XEN cell lines are derived not only from PrE of blastocysts but also from epiblasts. By deriving post-XEN cells from clumps of epiblasts and PE and VE and deriving pre-XEN cell lines from single PrE and single epiblasts of blastocysts.

In the process of deriving pre-XEN cell lines from blastocysts, the success rate in ES medium with LIF (56%) is higher than in TS cell medium with F4H (21%)^31^. ES medium with LIF is typically used to derive ES cell lines and in parallel XEN cell lines, and TS cell medium with F4H to derive TS cell lines and in parallel XEN cell lines^33^. The reason of the derivation pre-XEN cell lines in ES medium with LIF higher than in TS medium with F4H, could be LIF promote epiblast (ES-like) cells growth, and then enough epiblast (ES-like) cells convert to XEN cells. LIF had been reported to promote PrE cells in blastocysts^34^ and could also be promote XEN cells. Thus, it could be pre-XEN cells derived in ES medium with LIF have more cells originally from epiblast, and then improve the success rate of pre-XEN cell lines, and it further supports our conclusion that pre-XEN cells are both from PrE and epiblast. For this reason, we derived pre-XEN cell lines from ICMs with high efficiency (∼90%, 24 pre-XEN cell lines from 27 ICMs) that cultured the ICMs in ES medium with LIF, or in ES medium with LIF for 7 days, and then replaced the medium by TS medium with F4H (data not shown).

We never found flat colonies in the culture by derivation of post-XEN cell lines from the whole embryo method of E5.5 and E6.5^19^. However, the flat colonies were always cultured by the disaggregated embryo method on the earlier days and could not keep the flat colonies after passaging the cells by disaggregating the cells to single cells by TryPLE Express. Immunofluorescence showed that OCT4 and NANOG were expressed in flat colonies on day 18, and GATA6 and SOX7 expression surrounded flat colonies by the disaggregation method (Fig. 1G). OCT4 and Nanog are expressed in the epiblasts of E6.5 embryos^35^, and by the disaggregated embryo method, epiblast cells formed flat colonies with OCT4 and NANGO expression, and then the flat colonies gradually converted to XEN cells that surround the flat colonies. By the whole embryo method, epiblasts of E6.5 embryos mostly lost OCT4 and NANGO expression, did not form flat colonies, and could be directly converted into XEN cells. Outgrowths from single epiblasts of blastocysts are very similar to flat colonies from the disaggregated embryos method. Outgrowths from single epiblasts of blastocysts expressed OCT4 in the middle of the flat clone, and GATA6 surrounded the flat clone on day 7. After passaging the outgrowths from single epiblasts of blastocysts and culturing on day 15, the flat colonies both expressed OCT4 and NANOG and weakly expressed GATA6 and SOX7 (Fig. 4E), which is similar to the flat colonies from E6.5 embryos obtained by the disaggregation method. It could be that epiblast-derived cells can convert more easily to XEN cells in the whole-embryo method than in the disaggregated-embryo method.

Here, we revealed a mixed origin of XEN cell lines. How can epiblasts of blastocysts and postimplantation embryos contribute to XEN cell lines? ICM comprises three distinct cell types. The first type coexpresses NANOG and GATA6 and differentiates into epiblasts or PrE; the second type forms epiblasts; and the third type forms PrE^1^. ES cells go from an epiblast-like stage to a NANOG+ GATA6+ coexpression stage, over to a PrE-like stage, and then convert into XEN cells^36^. We speculate that, likewise, cells from a blastocyst could go from the NANOG+ GATA6-epiblast stage to the NANOG+ GATA6+ coexpression stage, over to a NANOG-GATA6+ PrE stage, and then contribute to a pre-XEN cell line. Similarly, epiblasts of postimplantation embryos could pass through a NANOG+ GATA6+ stage to a NANGO-GATA6+ stage and contribute to a post-XEN cell line. Epiblast cells of postimplantation embryos are differentiated from epiblasts of blastocysts, so they could be much easier to convert to XEN cell lines than ES cells.

We have previously reported the derivation of post-XEN cell lines from E5.5 and E6.5 embryos^19,20^; we were not able to derive post-XEN cell lines from E7.5 embryos using the whole-embryo method. Here, we derived PE-XEN cell lines and VE-XEN cell lines from E7.5 embryos using the disaggregation method. We have thus shown that PE and VE each can give rise to post-XEN cell lines and that E7.5 embryos can give rise to post-XEN cell lines. VE cells transdifferentiate into PE cells, and PE cells mimic the characteristics of XEN cells^28,37^.

It will be interesting to follow the transition of cells during the course of establishing a pre-XEN or post-XEN cell line from PrE and derived cell ExEn, and epiblast of blastocyst and postimplantation embryo, for instance by videomicroscopy, and to subclone XEN cell lines derived from (R26-tauGFP41 x Sox2-Cre) F1 embryos, and the process of epigenetic modification, in order to search for possible differences between PrE/ExEn-derived and epiblast-derived XEN cells.

## METHODS

### Mouse strains

The R26-tauGFP41 reporter strain ^22^ was a gift from Dr. Uli Boehm, Universität des Saarlandes, Homburg, Germany. Sox17-Cre strain^23^ was obtained from MMRRC, strain 036463-UNC, strain name Sox17<tm2(EGFP/cre)Mgn>/Mmnc. Although this strain contains *GFP*, and we can confirm the presence of *GFP* in the targeted insertion in the *Sox17* locus, we cannot detect GFP expression in embryos and cell lines. The Sox2-Cre strain^27^ was obtained from The Jackson Laboratory, strain #14094, strain name B6N.Cg-Tg(Sox2-cre)1Amc/J. Mouse embryonic fibroblasts (MEFs) were prepared using mitomycin C at E13.5 from strain #3208, strain name Tg(DR4)1Jae, obtained from The Jackson Laboratory.

### Cell culture

For TS cell medium, advanced RPMI-1640 (Gibco #12633-012) was supplemented with 20% (vol/vol) FBS (HyClone #SH30071.03), 2 mM GlutaMAX Supplement (Gibco #35050), 1% penicillin/streptomycin (Specialty Media #TMS-AB2-C), 0.1 mM β-mercaptoethanol (Gibco #21985-023), 1 mM sodium pyruvate (Gibco #11360-039), 25 ng/ml FGF4 (Peprotech #100-31) and 1 *µ*g/ml heparin (Sigma #H3149). For ES cell medium, DMEM (Specialty Media #SLM-220) was supplemented with 15% FBS (HyClone #SH30071.03), 2 mM GlutaMAX, 1% penicillin/streptomycin, 1% β-mercaptoethanol (Specialty Media #ES-007-E), 0.1 mM nonessential amino acids (Gibco #11140-035), 1 mM sodium pyruvate, and 1000 IU/ml leukemia inhibitory factor (LIF) (Millipore #ESG1107). Wells were coated with gelatin (Specialty Media #ES-006-B) and covered with MEF.

### Immunofluorescence and imaging

Cells were cultured in 4- or 24-well dishes. Cells were fixed in 4% paraformaldehyde at 4°C overnight or at room temperature for 30 min, permeabilized with 0.1% Triton X-100 in 1× PBS (1× PBST) for 30 min, and blocked for 1 hr with 5% normal donkey serum (Jackson ImmunoResearch Laboratories #017-000-121) diluted in 1× PBST (blocking solution). Primary antibodies were diluted at 1:50-1:500 in blocking solution, and samples were incubated at 4°C overnight. After three 10-min washes in 1× PBST, samples were incubated for 1-1.5 hr at room temperature in a 1:500 dilution of secondary antibody in blocking solution, washed and covered with 1× PBST containing DAPI. Primary antibodies from Santa Cruz Biotechnology were against GATA4 (#SC-1237), DAB2 (#SC-13982), OCT3-4 (#SC-5279), NANOG (#SC-376915), PDGFRA (#SC-338), and CDX2 (#SC-166830). Primary antibodies from R&D Systems were against GATA6 (#AF1700), SOX7 (#AF2766), and SOX17 (#AF1924). Primary antibodies against E-cadherin (ECCD2) were purchased from Invitrogen (#13-1900). Secondary antibodies from Jackson ImmunoResearch Laboratories were Cy5 AffiniPure Donkey anti-goat IgG (H+L) (#705-175-147). Secondary antibodies from Invitrogen were donkey anti-rabbit IgG (H+L) with Alexa Fluor 546 (#A10040) and donkey anti-mouse IgG with Alexa Fluor 546 (#A10036).

## Ethics statement

All animal studies were carried out in accordance with the German Animal Welfare Act, European Communities Council Directive 2010/63/EU, and institutional ethical and animal welfare guidelines of the Max Planck Institute of Biophysics and the Max Planck Research Unit for Neurogenetics. All experimental protocols were approved by the *Regierungspräsidium* Darmstadt and the *Veterinäramt* of the City of Frankfurt.

## Data availability

The datasets generated and/or analyzed during the current study are available from the corresponding author on reasonable request.

## Acknowledgments

J.L. is grateful to the Max Planck Society for generous financial support.

## Author Contributions

J.L. designed the research, performed the experiments, analyzed the data and wrote the manuscript.

## Additional Information

### Competing financial interests

The authors declare no competing financial interests.

## Notes

### Competing Interest Statement

The authors have declared no competing interest.

